# Source-space EEG functional connectivity and prediction of cognition in Parkinson’s disease: No added benefit of individualized head models over standard templates

**DOI:** 10.64898/2026.05.07.723671

**Authors:** Alina Tetereva, Grace Hall-McMaster, Nicola Slater, Abbie Harris, Reza Shoorangiz, Campbell Le Heron, Ross Keenan, Daniel Myall, Toni Pitcher, Ian Kirk, Wassilios G. Meissner, Tim Anderson, Tracy Melzer, Narun Pat, John Dalrymple-Alford

**Affiliations:** New Zealand Brain Research Institute, Christchurch, New Zealand; Te Kura Mahi ā-Hirikapo | School of Psychology, Speech and Hearing, University of Canterbury, Christchurch, New Zealand; University of Otago, Dunedin, New Zealand; Department of Medicine, University of Otago, Christchurch, New Zealand; Department of Neurology, Christchurch Hospital, Te Whatu Ora Waitaha Canterbury, Christchurch, New Zealand; RHCNZ Medical Imaging Group, Christchurch, New Zealand; School of Psychology, University of Auckland, Auckland, New Zealand; Centre for Brain Research, University of Auckland, Auckland, New Zealand; CHU Bordeaux, Service de Neurologie des Maladies Neurodégénératives, IMNc, Bordeaux, France; Univ. de Bordeaux, CNRS, IMN, UMR 5293, Bordeaux, France

**Keywords:** Parkinson’s disease, cognition, electroencephalography, functional connectivity, source localisation, head modelling, machine learning

## Abstract

**Introduction:** Cognitive decline is a major non-motor feature of Parkinson’s disease (PD), but reliable and accessible biomarkers remain limited. Resting-state electroencephalography (EEG) is a promising candidate because it is low-cost, portable, and well suited to repeated assessment. Recent work has increasingly focused on source-space functional connectivity (FC) for the prediction of cognition. However, the influence of source modelling based on an individualized MRI-based head model relative to that based on standard template model is unknown.

**Methods:** To compare these two source-space EEG FC methods, we analysed EEG data from the New Zealand Parkinson’s Progression Programme, including 136 people with PD and 51 age-similar controls. Source space resting-state EEG, parcellated with the HCP-MMP1 atlas, was used to derive amplitude envelope correlation (AEC) and debiased weighted phase lag index (dwPLI) across six canonical frequency bands. The resulting twenty-four FC modalities were evaluated using six machine-learning regression algorithms within a nested cross-validation framework.

**Results:** Theta-, alpha-, and beta-band FC showed the most consistent prediction of global cognition. The strongest performance was observed for theta- and alpha-band AEC and dwPLI features (max R² = 0.170, 95% CI = 0.067–0.262; max r = 0.439, 95% CI = 0.328–0.537). Standard and individualized head models showed comparable predictive performance across nearly all modalities. The feature-importance neuroanatomical patterns for Cole-Anticevic networks were also similar between the two head-model options.

**Conclusions:** We found that source-space resting-state EEG FC can predict cognitive performance in PD. The comparability of the two head models suggests that the more user-friendly and less resource-intensive standard template head model is sufficient for this purpose. This supports feasible, scalable, and clinically accessible EEG-based FC biomarkers of cognition in PD.

## Introduction

Parkinson’s disease (PD) is a progressive neurodegenerative disorder characterized by a variety of motor and non-motor symptoms (Schapira et al., 2017; Burchill et al., 2024). Cognitive decline is one of the most prominent non-motor symptoms, with a substantial proportion of patients experiencing a marked reduction in quality of life as well as progression to Parkinson’s disease dementia (PDD) (Lawson et al., 2016; Lawson et al., 2017; Gallagher et al., 2024). However, cognitive impairment in PD is highly heterogeneous, making early detection of cognitive deficits crucial for timely intervention and effective disease management (Aarsland et al., 2021; Weintraub et al., 2022). Despite ongoing efforts, reliable and accessible biomarkers capable of tracking cognitive decline in PD remain limited (Droby et al., 2022; Fu & Halliday, 2025; Guo et al., 2025).

Brain activity associated with cognitive decline can be directly and non-invasively detected using quantitative electroencephalography (qEEG) (Droby et al., 2022; Lee & Lee, 2025). Compared with magnetic resonance imaging (MRI) or positron emission tomography (PET), EEG offers high temporal resolution, is low-cost, more widely available, portable and generally acceptable to patients even as cognition worsens when examined using task-free (resting state) EEG (Babiloni et al., 2021; Choi et al., 2024). These advantages make EEG particularly suitable for assessing neural activity in clinical populations and in large-scale community studies (Abreu et al., 2020; Dierks et al., 2000).

An increasing number of studies have investigated EEG metrics in PD (Liao et al., 2024) and their association with cognition (Babiloni et al., 2018a; Betrouni et al., 2019; Chaturvedi et al., 2019; Sweeney et al., 2020; Geraedts et al., 2021; Parajuli et al., 2023; Mostile et al., 2025; Sasidharan et al., 2025). Researchers have begun moving towards source-space EEG and functional connectivity (FC) approaches to capture more anatomically and functionally meaningful neural connectivity (Anjum et al., 2024; Del Percio et al., 2025; Spoa et al., 2025). This shift reflects a broader view that cognitive impairment is increasingly understood as a disorder of large-scale brain networks. FC provides a window into the coordinated connectivity between brain regions, offering a more comprehensive characterization of brain function than regionally isolated measures (Abreu et al., 2020; Liu et al., 2018; Babiloni et al., 2018b). Connectivity in source space enhances insights by improving anatomical interpretability, approximating underlying cortical generators, and helping to reduce the impact of volume conduction (Liu et al., 2023; Vorwerk et al., 2014; Ramon et al., 2006; Hallez et al., 2008). In parallel, the integration of machine learning techniques with EEG-based connectivity measures have gained momentum. These approaches enable the extraction of complex high-dimensional patterns and the identification of latent network relationships, thereby improving the prediction of cognitive performance and potentially identifying individuals with PD at elevated risk for cognitive decline (Almgren et al., 2023; Anjum et al., 2024; Zawislak-Fornagiel et al., 2025).

The influence of preprocessing and modelling pipelines in this context, however, has not been systematically evaluated. The relative value of head modelling strategies for source reconstruction, the frequency bands analysed and different connectivity metrics have not been resolved (Bénar et al., 2002; Hallez et al., 2008; Ramon et al., 2006; Vanrumste et al., 2000; Vorwerk et al., 2014; Fuchs et al., 2001). EEG source imaging frameworks have recommended anatomically informed modelling to improve the localization of underlying neural generators (Liu et al., 2018; Michel et al., 2019). Head model selection can be based on either a standard head template or each individual’s MRI, which may differently influence the accuracy of source localization. Standard template models, however, may introduce greater uncertainty in source localization due to spatial mixing and imprecise modelling of individual head anatomy. This is important because inaccuracies at source reconstruction can propagate through subsequent analyses, potentially altering estimates of functional connectivity (Liu et al., 2023). Such differences may ultimately affect the performance and interpretability of machine learning models trained on these features.

We therefore evaluated the extent to which head model selection influences the robustness and reproducibility of EEG-based biomarkers of cognition. If individualized MRI-based head models do not substantially outperform standard template models, this would markedly enhance the clinical feasibility of EEG-based assessments of cognitive change. Standard head templates offer a more cost-effective, practical, and accessible alternative, whereas requiring individual MRI scans would substantially increase financial and resource demands. The current study addressed these questions in the context of predicting current cognition in PD. We directly compared the predictive performance of a standard head model relative to an individual MRI-based head model in the same participants. We compared both power-based and phase-based measures of FC across the range of canonical frequency bands, using the HCP-MMP1 parcellation and Cole-Anticevic network atlas as a means to extract brain networks across the entire cortex.

## Methods

### Participants

We used data from participants in the New Zealand Parkinson’s Progression Programme (NZP3; MacAskill et al., 2023) who provided both resting state (rs; “task-free”) EEG recordings and 3T structural MRI. Of the 194 participants, 187 remained after applying exclusion criteria (history of stroke [n = 1], brain tumours or cysts [n = 1], non-PD motor disease diagnosis [n = 3], and controls with cognitive impairment [n = 2]). Of these, 74 were female, and the mean age was M=71.5, SD=7.5 years. The sample comprised 136 individuals with PD and 51 similar-aged controls (Control). The study was approved by the Northern B Health and Disability Ethics Committee of the New Zealand Ministry of Health. At the beginning of their first neuropsychological assessment, all participants read and signed a consent form after discussing any questions or concerns. People with PD were diagnosed by consultant neurologists (TJA primarily) using Movement Disorder Society (MDS) criteria and were receiving prescribed medication at the time of all assessments.

Cognitive status was evaluated using the MDS Level II guidelines (Litvan et al., 2012; Emre et al., 2007), with neuropsychological testing across five cognitive domains (attention and working memory, executive function, visuospatial abilities, episodic memory, and language). Per Horne et al. (2021), the language domain was excluded from further analysis because it was not scored using external normative data. Scoring of other neuropsychological tests was conducted using age- and education-adjusted normative data (including sex-adjustment when available), with cognitive status and exclusion criteria are detailed in our previous studies (Horne et al., 2021). Neuropsychological outcomes alongside significant-other evaluation of activities of daily living resulted in 53 PD participants who were classified with mild cognitive impairment (PD-MCI). This was operationalised as two impairments at −1.5SD on two tests within any cognitive domain and everyday cognitive function not impaired, which tracks high risk of future PD with dementia (PDD; Wood et al., 2016). Current PDD was identified in 14 participants, based on impaired everyday cognition and deficits on at least one test at −2.0 SD in each of two domains. The remaining 69 PD participants were designated as showing normal cognition (PD-N). None of the control participants met criteria for MCI or had any history of neurological impairment. For our primary analyses, we computed a global cognitive score (global Z) by taking the mean of the average standardized scores within each of the four cognitive domains listed above.

### MRI Acquisition and Preprocessing

Structural MRI data were acquired on a 3T Siemens Skyra scanner (Pacific Radiology, St. George’s Hospital, Christchurch) equipped with a 64-channel head and neck coil. High-resolution T1-weighted images were obtained using a standard MPRAGE sequence (TR = 2000 ms, TE = 2.85 ms, TI = 880 ms, flip angle = 8°, field of view = 256 mm, acquisition matrix = 256 × 256). The sequence produced 208 sagittal slices with an isotropic voxel size of 1 × 1 × 1 mm³ (bandwidth: 240 Hz/pixel). All T1-weighted images were preprocessed using FreeSurfer version 7.2 (Fischl, 2012). Cortical and subcortical reconstruction and segmentation were performed with the recon-all pipeline, which includes intensity normalization, skull stripping, surface tessellation, topology correction, and surface reconstruction. The outputs were visually inspected to ensure accurate cortical surface delineation.

### EEG Recording Procedure

Resting-state EEG was collected using a 64-channel Quick-Cap Neo Net system (Compumedics Neuroscan) with Ag/AgCl electrodes. Data were acquired through a SynAmps2 amplifier and recorded in Curry 7 software (Compumedics Neuroscan) at 24-bit resolution. The reference electrode was placed midway between Cz and CPz, and AFz served as the ground. Signals were sampled at 500 Hz to capture the relevant frequency range and minimize aliasing (Mahajan et al., 2025). A vertical EOG channel was included to assist in detecting eye movement–related artifacts. Electrode impedances were kept below 10 kΩ for the duration of the recording.

Each participant completed a continuous resting-state session of approximately 10 minutes. The session consisted of three eyes-closed blocks of about 3 minutes each, separated by brief breaks during which participants were reminded to stay awake. This block structure maintained alertness while reducing state fluctuations during the recording.

### EEG preprocessing pipeline

EEG preprocessing was conducted using a modular approach that combined the steps from a standardized pipeline MNE-BIDS-Pipeline (v1.9) (Appelhoff et al., 2019; https://mne.tools/mne-bids-pipeline/1.9/features/steps.html; https://github.com/mne-tools/mne-bids-pipeline/tree/main/mne_bids_pipeline/steps/preprocessing) with our custom Python scripts (see https://github.com/HAM-lab-Otago-University/NZP3_Chch_cog_pred_STDvsIND_headmodels). Some preparatory steps were carried out manually or with additional packages outside the pipeline. Continuous EEG recordings were first cropped to retain only the period between the experiment’s designated start and end markers. Bad channels were identified using visual inspection, and the NoisyChannels algorithm from the pyPrep package (Bigdely-Shamlo et al., 2015), and their labels were saved into subject-specific TSV files. To standardize the data across participants, some electrodes that were not consistently present due to minor differences in head cap configuration were removed (F11, F12, FT11, FT12, FT7, FT8, PO5, or PO6), yielding a final set of 58 common EEG channels. In addition, the two mastoid channels and the vertical electrooculogram (VEO) were excluded due to frequent signal dropout or noise. Channel names were updated to comply with BIDS conventions.

Bandpass filtering was applied from 0.5 to 100 Hz using zero-phase FIR filters (MNE-BIDS-Pipeline steps _01_data_quality and _04_frequency_filter). All recordings were downsampled to 250 Hz to reduce computational load, and EEG signals were re-referenced to the average reference. A separate custom script was used to remove power line interference. Powerline noise centered at 50 Hz was reduced using the “Denoising Source Separation” (DSS) method from the meegkit package (meegkit.dss.dss_line) (de Cheveigné & Simon, 2008), with six DSS components removed from each dataset to attenuate narrowband contamination.

Independent Component Analysis (ICA) was applied using the Picard algorithm with extended-infomax settings, via pipeline steps (MNE-BIDS-Pipeline steps _06a1_fit_ica and _06a2_find_ica_artifacts). Initial artifact components were flagged automatically using autoreject_local. A separate custom script was used to automatically label ICA components using the mne_icalabel package, which implements the ICLabel classifier (Pion-Tonachini et al., 2019). Components classified as non-brain sources (e.g., ocular, cardiac, muscle, line noise) were marked for exclusion, and these labelled components were removed from the data during later pipeline step (MNE-BIDS-Pipeline step _08a_apply_ica from MNE pipeline).

The cleaned EEG signal was segmented into 3-second epochs that had 1.5-second overlap (during the MNE-BIDS-Pipeline step _07_make_epochs). Epoch-level artifact rejection was then performed using the autoreject package (MNE-BIDS-Pipeline step _09_ptp_reject), which detects outlier channels based on channel-wise peak-to-peak amplitude thresholds. Within each epoch, channels exceeding the artifact threshold were interpolated when possible (up to 32 channels per epoch), and entire epochs were discarded when the level of distortion was higher than the max threshold. Epochs occurring outside of the designated three resting-state blocks were excluded from analysis using a custom script (marked as bad epochs).

Bad channels previously identified were interpolated using spherical spline interpolation (mne.io.Raw.interpolate_bads) in a custom script, applied separately to both continuous and epoched data. A predefined list of unreliable channels, including mastoids and VEO, was excluded prior to interpolation. After interpolation, signals were again re-referenced using projection mode. A diagonal noise covariance matrix was computed from the cleaned raw data using the empirical method in MNE-Python (mne.compute_raw_covariance followed by .as_diag()), and used for inverse modelling to generate source localisation.

### Head modelling and source localisation

Boundary element method (BEM) surfaces were constructed using FreeSurfer’s (Fischl, 2012) watershed algorithm for both the standard (template) head model (*fsaverage*) and each participant’s individual anatomical MRI (individual head model). Coregistration between EEG sensor positions and MRI anatomy was performed using fiducial alignment and iterative closest point (ICP) fitting, with additional constraints applied to the nasion, preauricular points, and digitized scalp surface. Minor manual rotation adjustments were applied to optimize alignment. The same coregistration procedure was used for both head model types; for the standard template, this was performed once and then applied to all participants, whereas for individual head models, coregistration was performed separately for each participant’s MRI. The resulting transformation matrices were saved for subsequent source modeling. Three-layer BEM models were created using standard conductivity values [0.3,0.006,0.3] for the scalp, skull, and brain. Source spaces were defined using oct6 spacing, and forward solutions were computed using the standard template (fsaverage template) (Fischl et al., 1999) and individual (i.e. subject-specific) head models and coregistration transforms.

Prior to source projection, epoched EEG data were re-referenced to the average reference. Source estimates were calculated using minimum norm estimation, with a regularization parameter *λ*^2^ =1/9 and dipole orientations constrained to the cortical surface normal. Both standard and individual head models were used to generate source localisation and then analyses based on each were generated and compared. Source estimates were parcellated into 360 cortical regions using the HCP-MMP1 atlas (Glasser et al., 2016), and regional-averaged time series were extracted using the mean_flip method. These procedures were performed separately for spectral analysis for each of the six canonical frequency bands: delta (0.5–4 Hz), theta (4.01–7.99 Hz), alpha (8–12.99 Hz), beta (13–30 Hz), gamma1 (30.01–49 Hz), and gamma2 (51–80 Hz).

### EEG functional connectivity calculation

We computed two FC metrics using 360 regionally averaged source-space time series from the HCP-MMP1 atlas, with connectivity matrices generated for each subject, session, frequency band, and head model. First, power-based FC was computed using the amplitude envelope correlation (AEC), estimated by band-pass filtering the atlas-extracted source-space time series within each frequency range, followed by Hilbert transformation to obtain analytic envelopes. Pairwise orthogonalization was applied to reduce spurious correlations due to signal leakage, and AEC was computed using the envelope_correlation function from the mne_connectivity package (MNE-Connectivity Developers, 2024). Second, phase-based FC was computed using the debiased weighted Phase Lag Index (dwPLI), applied to the same atlas-extracted time series. The spectral_connectivity_epochs function from mne_connectivity was used with Fourier-based decomposition and frequency averaging enabled.

We additionally assigned each of 360 regions to large-scale resting-state functional networks using the Cole–Anticevic network atlas (Ji et al., 2019), which is derived from the same HCP-MMP1 parcellation. This atlas groups regions into 12 canonical networks: Visual1, Visual2, Somatomotor, Cingulo-Opercular, Language, Default, Frontoparietal, Auditory, Posterior-Multimodal, Dorsal-attention, Ventral-Multimodal, Orbito-Affective. Each of the 360 regions was labelled according to its network membership, enabling subsequent analyses of within- and between-network functional connectivity patterns.

In total, we derived 24 modalities, representing all combinations of the two FC metrics, two head models, and six frequency bands. Thus each modality corresponded to a unique combination of these factors, such as AEC estimated using the standard head model in the alpha band, or dwPLI estimated using the individual head model in the theta band. For each subject and modality, the connectivity matrix comprised 64,620 unique pairwise connections between 360 brain regions (lower triangle of the 360 × 360 matrix, excluding the main diagonal). These matrices were vectorised by flattening the lower triangle into a one-dimensional feature vector and then assembled across subjects, yielding one table per modality with dimensions N × 64,620, where N is the number of subjects. Each column in these tables represented a single brain feature, defined as one pairwise functional connection, or edge, between two brain regions.

### Machine Learning Pipeline

To evaluate the ability of each modality to predict cognitive performance, and minimize model over-fitting, we implemented a nested cross-validation framework using scikit-learn (Pedregosa et al., 2011). Consistent with our earlier work (Tetereva et al., 2022; 2025), the nested cross-validation comprised an outer loop providing unbiased estimates of predictive performance and an inner loop for hyperparameter tuning. For the present study, we split the dataset into four outer folds using the KFold function, ensuring that each fold maintained a comparable ratio of participants with Parkinson’s disease (PD) and Control. In each outer iteration, one fold served as the held-out test set (∼25% of the data) whereas the remaining folds (∼75%) were used for model training and hyperparameter tuning via grid-search cross-validation in the inner loop. This procedure ensured that model selection was performed exclusively within the training data and that no information from the outer test set contributed to hyperparameter optimisation, thereby preventing data leakage. In each outer-fold iteration, the held-out test set was not used for model selection, hyperparameter optimisation, standardisation, target transformation, or any other model-fitting step. Within each outer-fold iteration, each modality (that is, from one of two FC measures × two head models × six spectral frequencies) and the target variable (global Z) were standardised independently (column-wise), with parameters estimated from the outer-fold training set only and subsequently applied to the corresponding test set. For all models, we implemented a unified preprocessing pipeline in which predictors were standardised using StandardScaler, while the target variable was modelled through a TransformedTargetRegressor that applied its own internal pipeline consisting of standardisation followed by a Yeo–Johnson power transformation. No residualisation or covariate adjustment was applied to either the EEG connectivity features or the target cognitive score. The target variable was derived from neuropsychological test scores that had already been standardised using available normative data adjusted for age and education, and for sex when available. The same preprocessing strategy was applied consistently across all 24 modalities.

We fitted six multivariate regression machine-learning algorithms using scikit-learn. These were: Elastic Net (Enet), Partial Least Squares (PLS) regression, Kernel Ridge Regression (KRR), Support Vector Regression (SVR), Random Forest (RF), and XGBoost (XGB). Elastic Net (Zou & Hastie, 2005) was used as a linear, regularised model combining Lasso (L1; coefficient elimination) and Ridge (L2; multicollinearity) penalties and is well suited for high-dimensional, correlated feature spaces; we tuned the regularisation strength α over 70 logarithmically spaced values between 10⁻¹ and 10² and the l₁-ratio over 25 values between 0 and 1. PLS regression (Wold et al., 2001), a linear dimensionality-reduction approach that extracts latent components maximising covariance between predictors and the target, was tuned over 1–200 components. KRR (Saunders et al., 1998) extends Ridge regression with nonlinear feature mapping through kernel functions, enabling it to capture smooth nonlinear relationships; we evaluated both linear and radial basis function (RBF) kernels, tuning α between 10⁻³ and 10³ and, for RBF kernels, γ between 10⁻³ and 10². SVR (Drucker et al., 1996), a margin-based kernel regression method capable of modelling nonlinear interactions while controlling model complexity, was tuned by varying C between 10⁻² and 10², γ between 10⁻³ and 10¹, and comparing linear and RBF kernels. Random Forest (Breiman, 2001), a tree-based ensemble method that captures nonlinearities and high-order interactions, was tuned over the number of trees (100–5000), maximum depth (None or 10–50), minimum samples per split (2–10), minimum samples per leaf (1–4), and feature-sampling strategy (sqrt, log2). XGBoost (Chen & Guestrin, 2016), a gradient-boosted tree algorithm that builds sequential trees to minimise residual error, was tuned over the number of estimators (100–500), maximum depth (1–9), learning rate (0.01–0.2), γ (0–5), and subsampling ratio (0.5–1.0). Hyperparameter tuning for all algorithms was performed using three-fold inner cross-validation, with the coefficient of determination (R²) used as the scoring metric. The best-performing configuration was then refit on the full outer-fold training set and evaluated on the corresponding held-out test set.

### Model Evaluation, Bootstrapping and Statistics

For each outer-fold, we computed predicted values of the corresponding held-out test set and compared them with the observed cognitive scores. Predictive performance was quantified using two complementary metrics: the coefficient of determination (R²), calculated as the sum-of-squares definition (R² = 1 - (Sum of Squared Residuals / Total Sum of Squares)), and Pearson’s correlation coefficient (r) for the linear association between predicted and observed values. To quantify uncertainty around these estimates, we aggregated predicted and observed values across all outer folds and generated 5,000 bootstrap samples (Efron & Tibshirani, 1993). For each bootstrap iteration, we recomputed R² and r yielding bootstrapped distributions and corresponding 95% confidence intervals for each modality and algorithm. To statistically compare model performance across modalities, we also calculated bootstrapped differences in each metric by subtracting the performance of one modality from another in every bootstrap sample. If the 95% confidence interval of this difference did not include zero, we concluded that the two modalities differed significantly in predictive performance.

To quantify the extent to which EEG-based biomarkers overlapped with age and sex in predicting cognition, we conducted a commonality analysis (Nimon et al., 2008). We focused on the three best-performing EEG modalities and performed the analyses on held-out test sets across all outer folds. Specifically, observed cognition was regressed on EEG-derived predicted cognition from each modality, along with age and sex. This approach enabled us to partition the explained variance, quantifying the unique contributions of EEG-based predictions as well as their shared effects with age and sex in predicting cognitive outcomes.

Additionally, feature importance (i.e., the relative contribution of individual brain features to model predictions) was estimated using a consistent approach, enabling comparability across models despite the use of multiple machine-learning algorithms. Feature importance was estimated using Haufe transformation (Haufe et al., 2014) by computing the Pearson correlation between each brain feature and the corresponding model-predicted values. This approach provides a consistent measure of feature relevance across algorithms without requiring extraction or transformation of model coefficients.

This procedure was applied to all features within each modality (two FC types × two head models x six frequency bands). Within each modality, feature importance values (correlation value) were rounded to three decimal places, converted to absolute values, and ranked from strongest to weakest. These ranked values were then visualised as half-matrix representations. To assess whether model predictions preserved group-level differences in cognitive status, we compared observed and predicted values across PD-N, PD-MCI, PDD and non-PD controls. Group differences were first evaluated using Welch’s ANOVA to account for unequal variances across groups, followed by Games–Howell post hoc tests for pairwise comparisons, for which Hedges’ g effect size was calculated. These analyses were applied to both observed and model-predicted values for each modality to determine whether predicted scores reflected the existing clinically meaningful group differences.

## Results

### Model performance across EEG functional connectivity modalities

Figure 1 shows the performance of the twenty-four modalities in terms of either AEC or dwPLI source-space FC metrics, standard head model (STD) or individual T1-derived head model (IND) and one of the six canonical frequency bands. The six multivariate regression machine-learning algorithms were applied to each of these modalities. Figure 1A shows the results for the best-performing algorithm within each modality (with maximum R² and Pearson’s r values indicated alongside the plots). For each modality, the numerically highest-performing algorithm was selected for presentation purposes, as statistically significant differences among the six machine-learning algorithms were not consistently observed (Figure S1). Supplementary Figure S1 compares the top-performing algorithm against the remaining algorithms for each modality via bootstrapping. In most cases, the top-performing algorithm did not differ significantly from the others.

**Figure 1.**
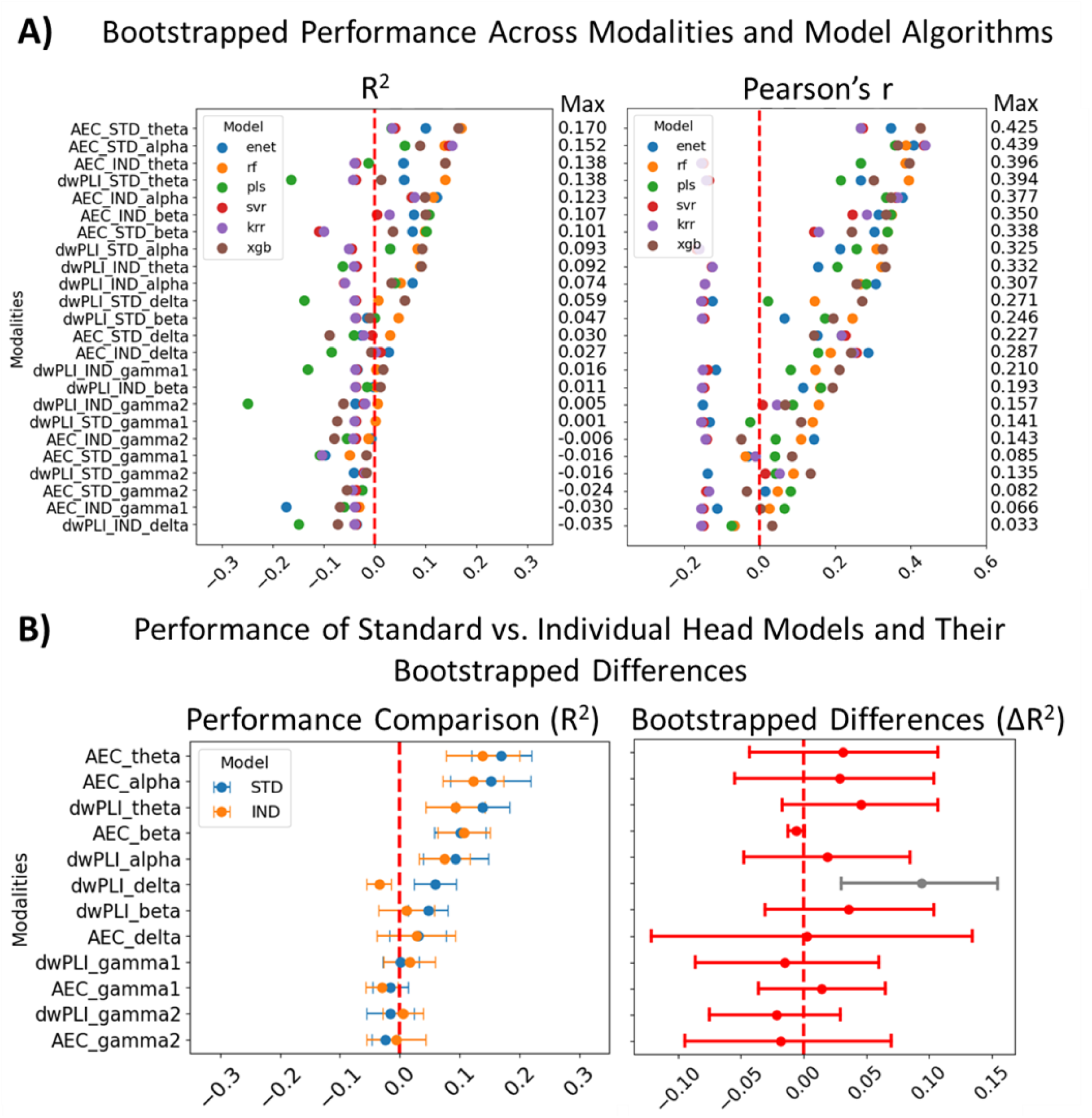
**A)** Bootstrapped performance for each machine-learning (ML) algorithm and 24 modalities, that is, two functional connectivity measures after source localisation derived from either the standard head model (STD) or the individual head models (IND) across six EEG frequency bands. Each coloured dot represents the bootstrapped mean estimate for each machine learning algorithm for each modality. Both the model coefficient of determination (R²) and Pearson correlation (r) for observed relative to predicted global Z are shown. The maximum performance value achieved across the six ML algorithms is shown on the right-hand side of each plot. The best prediction models are generally derived from theta and alpha frequency bands. **B)** Direct comparison of the STD versus IND head models’ performance, based on R². On the left, each modality (FC x frequency) is represented by two dots corresponding to the STD and IND head models. Each dot reflects the maximum model performance (R²) achieved across all algorithms for that modality (corresponding to the values shown in panel A). Dots represent bootstrapped means, and bars indicate standard deviation. On the right, the bootstrapped performance difference (ΔR²) between STD and IND models is shown for each modality. Dots represent mean differences, and bars represent 95% confidence intervals (CIs). Note, only one FC x frequency feature shows a difference (Grey bar; CI does not cross zero) between the models; red indicates no significant difference. AEC = amplitude-envelope; dwPLI = debiased weighted phase lag index.

Across the 24 modalities, FC in the theta and alpha bands consistently showed the strongest predictive performance for cognitive outcomes. For example, the top-performing modality AEC_STD_theta (RF) achieved R² = 0.170 (95% CI: 0.067–0.262) and r = 0.425 (95% CI: 0.304–0.537), while the next best-performing model, AEC_STD_alpha (KRR), achieved R² = 0.152 (95% CI: 0.009–0.272) and r = 0.439 (95% CI: 0.328–0.537); both of these modality performances were achieved using the standard template model. Corresponding individual head model modalities showed slightly lower performance, with AEC_IND_theta (XGB) achieving R² = 0.138 (95% CI: 0.011–0.256) and r = 0.396 (95% CI: 0.268–0.514), and AEC_IND_alpha (Enet) achieving R² = 0.123 (95% CI: 0.015–0.218) and r = 0.377 (95% CI: 0.257–0.483); however, differences between standard and individual head models were not statistically significant (Figure S2). These correlation values fall within the medium-to-large range based on contemporary benchmarks for effect size interpretation (Funder & Ozer, 2017; Gignac & Szodorai, 2016). Beta-band modalities also showed consistent predictive performance, although with numerically lower R² values than theta and alpha (e.g., AEC_STD_beta (PLS): R² = 0.101, 95% CI: 0.014–0.184; r = 0.338, 95% CI: 0.209–0.458). In contrast, FC modalities from other frequency bands yielded R² values near zero or negative, indicating limited explanatory power for individual variability in cognition. Delta-band AEC and dwPLI models failed to converge in approximately half of the cross-validation folds; therefore, results based on delta-band should be interpreted with caution.

Within the well-performing theta and alpha bands, AEC connectivity yielded numerically higher prediction performance than dwPLI, although these differences did not reach statistical significance (Figure S2). Although AEC_STD_theta was the numerically highest-performing modality, its performance did not differ significantly from other high-performing modalities, including AEC (theta, alpha, and beta bands) and dwPLI (theta and alpha bands) across both standard and individual head models. In contrast, these modalities showed significantly higher performance than the remaining 14 modalities in Figure 1, which exhibited R² values near zero or negative (Figure S2).

Figure S3 presents the commonality analysis, showing the variance in observed cognitive scores explained uniquely and jointly by the three best-performing EEG modalities, age, and sex in three-predictor models. Across both standard and individualized head models, EEG measures uniquely accounted for the majority of the variance in cognitive scores, with only minimal variance shared with age and sex.

### Comparison of standard vs. individual head models

Our findings suggest that, for the purpose of predicting cognitive performance from resting-state source-space EEG functional connectivity, individual T1 MRI scans for head modelling do not provide additional benefits. Figure 1B directly compares prediction performance between models built using the standard head model and the individual T1-based head model. No statistically significant differences were observed in the prediction of cognition between the standard head model and individual head model for any FC or frequency band. The exception was dwPLI in the delta frequency, but this should be interpreted with caution because delta frequencies often showed convergence failure. Standard head models yield comparable predictive performance and therefore provide a computationally efficient and clinically accessible alternative for source modelling in this context.

### Feature importance in standard vs individual head models

Feature importance was estimated as the Pearson correlation between each brain feature and the corresponding model-predicted cognitive scores and subsequently ranked within each modality.

Visual inspection of feature importance rank matrices also revealed minimal differences between standard and individual head models for the same modality (Figure 2), reiterating that the choice of head model had little impact on feature importance patterns in the prediction of cognition. This observation was supported by strong correlations between feature importance ranks derived from standard and individual head models across the best-performing modalities (Spearman ρ, all p < 0.001: AEC theta: ρ = 0.72, AEC alpha: ρ = 0.64, dwPLI theta: ρ = 0.44, AEC beta: ρ = 0.75, dwPLI alpha: ρ = 0.40), demonstrating consistent feature importance patterns across head models. Notably, AEC-based modalities showed higher correspondence between standard and individual head models (ρ = 0.64–0.75) compared to dwPLI-based modalities (ρ = 0.40–0.44), suggesting greater consistency of power-based connectivity patterns across head models.

**Figure 2.**
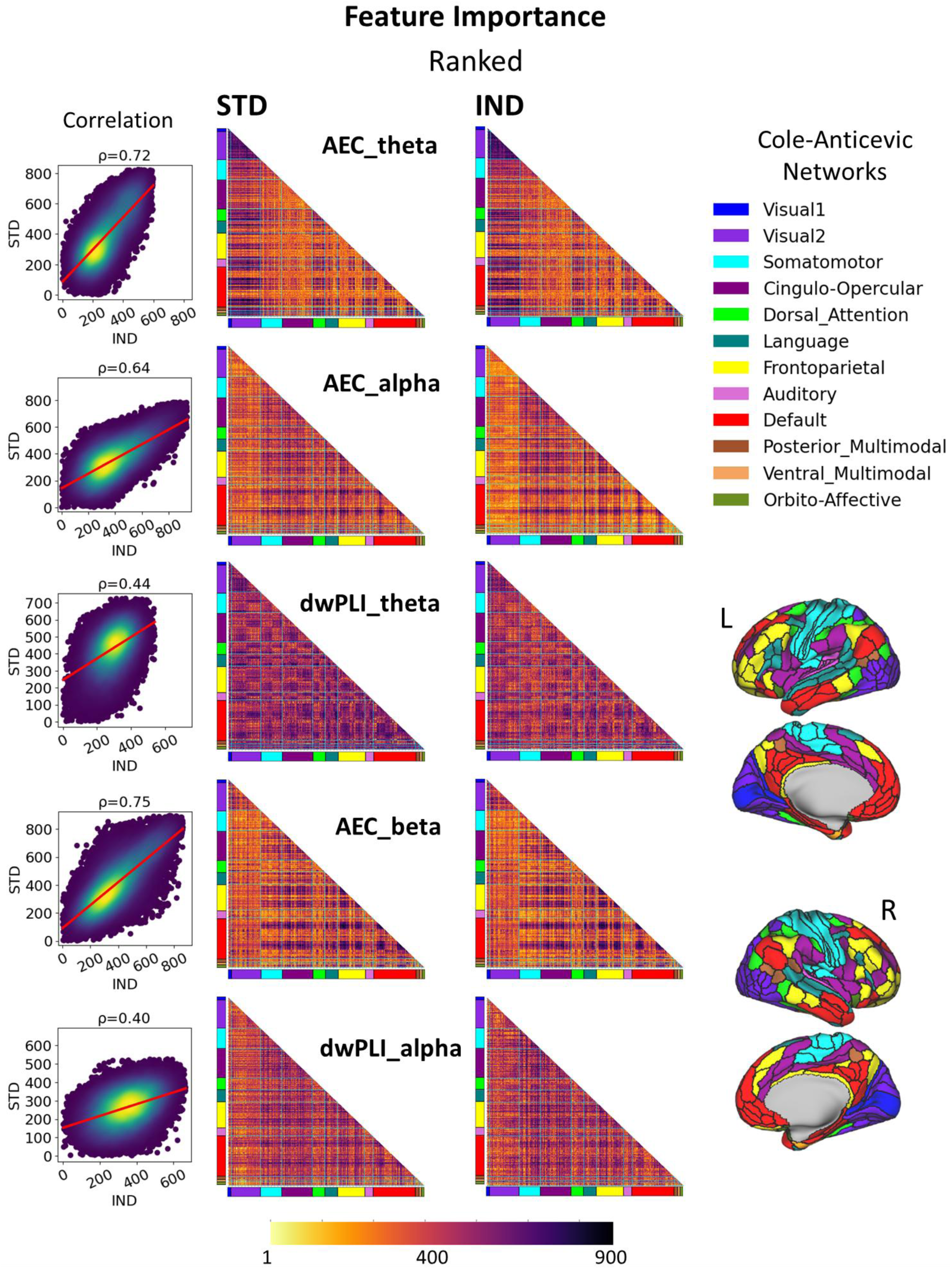
Comparison of feature importance patterns between standard and individual head models. Feature importance (FI) is shown for the five top-performing modalities (with their corresponding best-performing algorithms), comparing patterns between standard (STD) and individual (IND) head models. Feature importance was estimated as the Pearson correlation between each connectivity feature and the corresponding model-predicted values. Values were converted to absolute values and ranked within each modality from strongest to weakest. These ranked values are visualised as half-matrix representations, where stronger feature importance is shown in yellow and weaker (less informative) features are shown in darker colours. On the right, the Cole–Anticevic brain network parcellation is shown, including network labels and their cortical locations for reference. The left panel show scatter plots of FI ranks from STD and IND head models, with Spearman’s ρ used to quantify the similarity of feature-importance ranking patterns between the two head-model types.

To characterise the neuroanatomical patterns underlying model predictions, feature importance values were examined across large-scale functional networks defined by the Cole–Anticevic Brain Network Parcellation (Ji et al., 2019), which is based on regions from the HCP-MMP1 atlas (Glasser et al., 2016). Across the best-performing modalities, a clear directional pattern emerged. Theta-band connectivity was always negatively associated with cognitive performance, whereas alpha-band connectivity showedalways positive associations (Figure 3). For the AEC_theta modality, the most important features were primarily connections involving the Cingulo-Opercular and Somatomotor networks with other networks, as well as clusters linking Fronto-Parietal and Default Mode networks to the rest of the brain. In contrast, for AEC_alpha, a different pattern emerged, with prominent contributions from Visual2 network connectivity and clusters within the Default Mode network. Notably, Default Mode network connections showed a largely reversed feature importance pattern across these bands, with broadly high ranks in AEC_theta but low ranks in AEC_alpha, except for a few localized clusters. By contrast, no consistent between-network patterns were observed for dwPLI in either theta or alpha bands.

**Figure 3.**
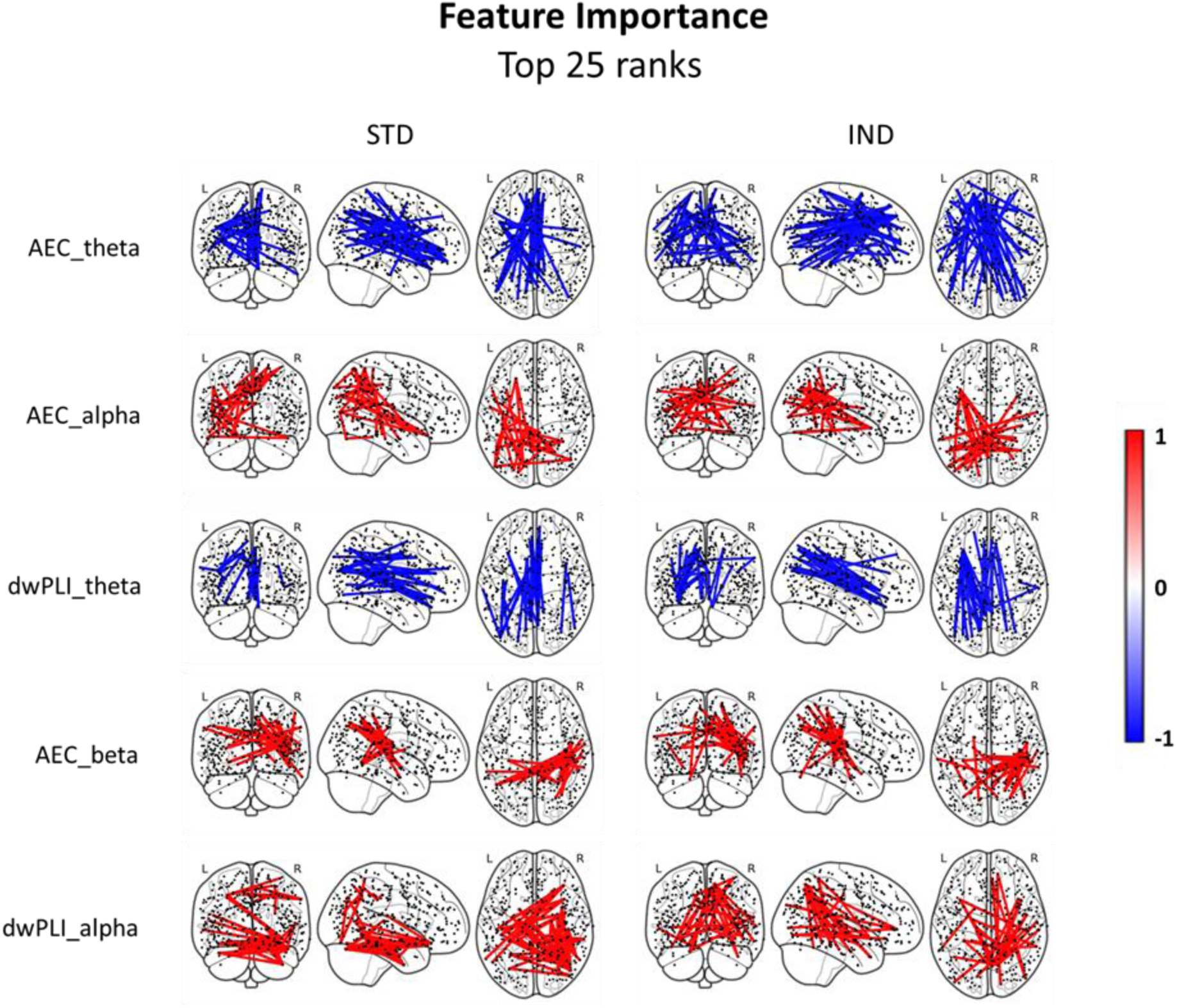
Spatial distribution of top-ranked feature importance values. Features corresponding to the top 25 FI ranks are shown. Because multiple features can share the same rank, this selection includes approximately 60–70 FC pairs per modality. Raw FI values are visualised and projected onto brain coordinates. FI was estimated as the Pearson correlation between each FC feature and the corresponding model-predicted values. Colour indicates the direction of the relationship between each feature and predicted cognition: red denotes a positive association (higher connectivity associated with higher cognitive scores), whereas blue denotes a negative association (lower connectivity associated with higher cognitive scores). The figure also allows comparison of spatial patterns between standard (STD) and individual (IND) head models across the same set of top-performing modalities.

Figure 3 illustrates these patterns by projecting the highest-ranked features onto brain coordinates, with standard and individual head models shown separately. Only the top 25 features are displayed (corresponding to approximately 50–70 FC pairs per modality) to balance interpretability and spatial coverage, as inclusion of a larger number of connections reduced clarity of network-level patterns. The spatial distribution and ranking of the top 25 features were highly similar between head models, consistent with the strong correspondence observed in Figure 2. Importantly, raw feature importance values (rather than ranks) are shown in Figure 3, allowing interpretation of the direction of associations between connectivity features and predicted cognition. The relationship between FC and cognition was always positive for the alpha band, whereas this association was always negative for the theta band. The opposing theta–alpha patterns indicate distinct large-scale network signatures associated with cognitive status.

### Group differences in observed and predicted cognitive scores

There was an expected group main effect for observed global cognition scores (F(3, 53.61) = 138.24, p < .001; Figure 4A). Post hoc tests showed that all group pairs differed significantly (all p < .01), with large effect sizes (Hedges’ g = 0.76–4.35), indicating clear separation between cognitive groups.

**Figure 4.**
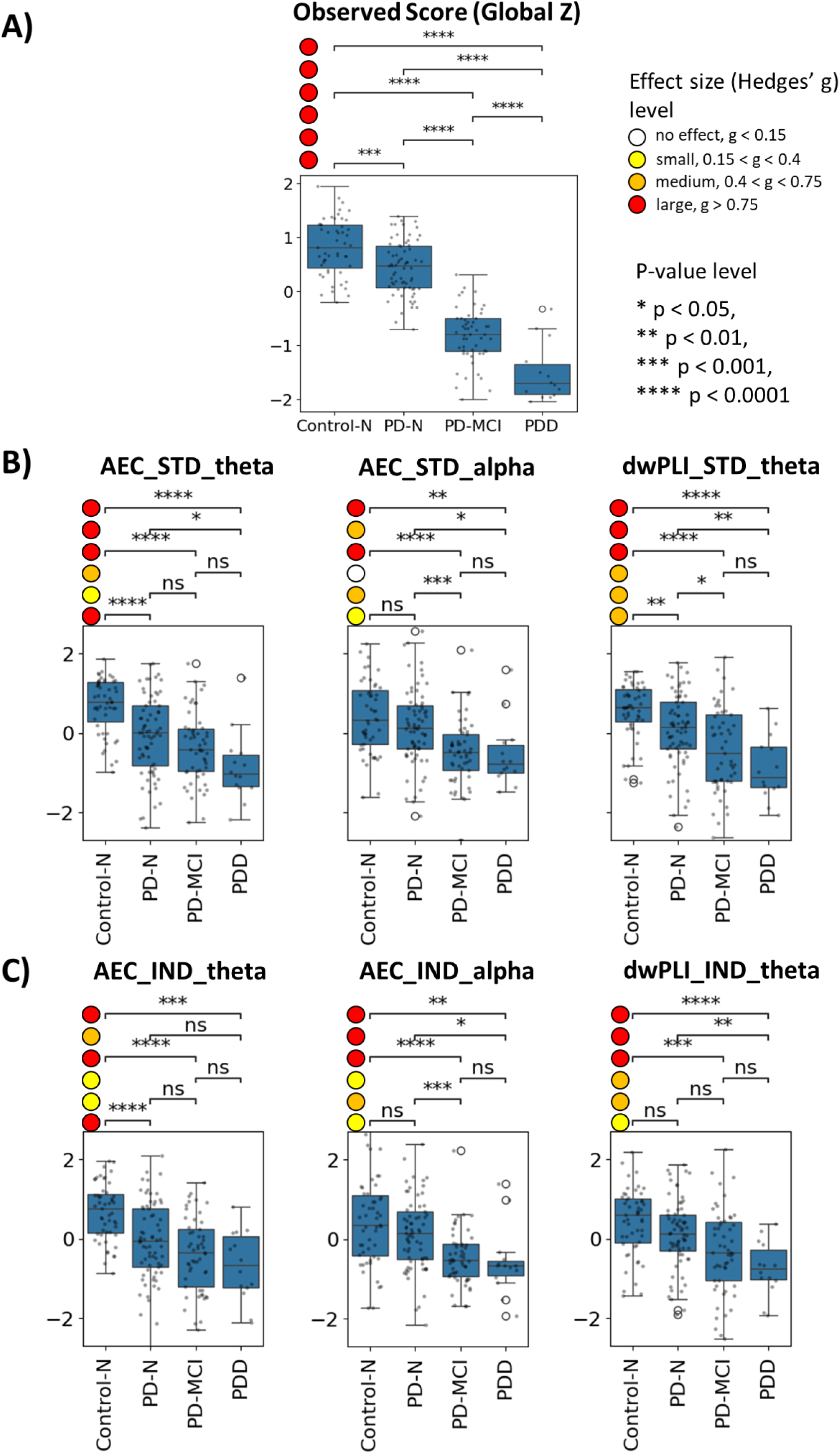
Group differences in observed and predicted cognitive scores. Panel A shows the distribution of observed global cognition scores (global Z). Panel B shows model-predicted scores for the three top-performing modalities based on standard head models (AEC_STD_theta, AEC_STD_alpha, and dwPLI_STD_theta), and Panel C shows the corresponding modalities based on individual head models (AEC_IND_theta, AEC_IND_alpha, and dwPLI_IND_theta). Brackets indicate pairwise group comparisons from Games–Howell post hoc tests, with asterisks denoting statistical significance. Circles shown to the left of each bracket indicate the corresponding Hedges’ g effect size (Brydges, 2019), with red indicating large effects, orange medium effects, yellow small effects, and white negligible effects.

Group main effects were also present in model-predicted values for the three best-performing standard head model modalities: AEC_STD_theta, AEC_STD_alpha, and dwPLI_STD_theta, with F(3, 54.32) = 22.78, p < .001, F(3, 55.43) = 11.31, p < .001, and F(3, 54.96) = 18.51, p < .001, respectively (Figure 4B). A similar pattern was observed for the corresponding individual head model modalities (AEC_IND_theta, AEC_IND_alpha, and dwPLI_IND_theta; F(3, 54.43) = 20.02, p < .001, F(3, 54.25) = 11.40, p < .001, and F(3, 58.43) = 11.56, p < .001, respectively; Figure 4C). The magnitude of the group main effects was broadly similar across standard and individual head model predictions, with theta-band modalities showing numerically larger F-values.

Compared with the observed cognitive scores, the predicted scores showed reduced differentiation between groups: all observed group differences were large, whereas predicted scores showed a mixture of small, medium, and large effects. Effect sizes were interpreted using thresholds proposed by Brydges (2019), with Hedges’ g values below 0.15 considered negligible, 0.15–0.40 small, 0.40–0.75 medium, and above 0.75 large. The largest effects in predicted scores were generally observed for comparisons involving more cognitively impaired groups, particularly Control-N versus PD-MCI, Control-N versus PDD, and PD-N versus PDD. Across these comparisons, effect-size patterns were broadly similar between predictions based on standard and individual head models, with no consistent evidence that individual head models produced stronger group separation. In contrast, differences between adjacent groups, such as PD-N versus PD-MCI and PD-MCI versus PDD, were less consistent.

Overall, these findings suggest that model-predicted values preserved the broad gradient of cognitive impairment across groups (Figure 5), but with reduced sensitivity to finer pairwise diagnostic distinctions compared with observed cognitive scores.

**Figure 5.**
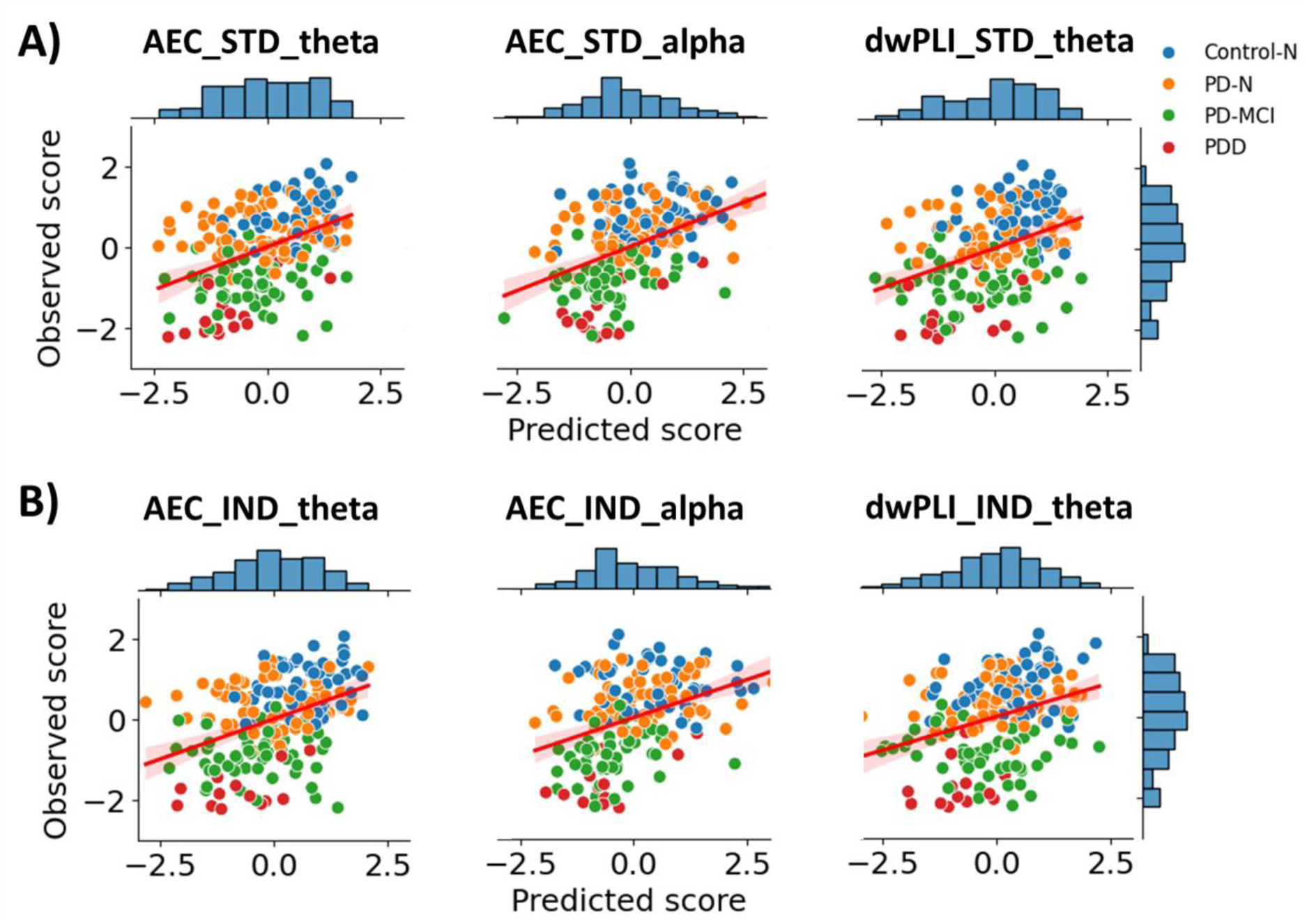
Observed versus predicted cognitive scores across diagnostic groups. The figure shows scatter plots of observed versus predicted cognitive scores (both standardised) for the three top-performing modalities. Panel A shows modalities based on standard head models, and panel B shows the corresponding modalities based on individual head models. Each point represents an individual participant, and colours indicate diagnostic groups. The observed global cognition scores (global Z) are shown on the y-axis, and model-predicted scores are shown on the x-axis. The plots demonstrate that predicted scores broadly preserve the relative ordering of individuals and partially retain group-level separation, with higher observed scores generally corresponding to higher predicted values. This is reflected in the positive association between observed and predicted cognition across modalities.

## Discussion

This study demonstrates that cognitive performance in people with PD can be predicted using source-space EEG FC. We found that individualized anatomical head models, based on 3T structural MRI, offered no advantage over standard template head models within a machine-learning framework for cognition-related EEG features in this cohort. Although further work is needed to determine whether this conclusion generalises to other neurological conditions, these results may have broader relevance, as the impact of head modelling on source reconstruction is primarily driven by general methodological and biophysical factors rather than disease-specific mechanisms.

Analysis across all twenty-four modalities revealed two additional key findings. First, EEG-based FC in the theta, alpha, and beta frequency bands consistently predicted cognitive performance across participants, whereas FC derived from delta and gamma frequencies showed weak or unstable predictions. Second, power-based (AEC) and phase-based (dwPLI) FC showed broadly comparable predictive performance, with numerically higher values observed for AEC in several top-performing modalities.

### EEG functional connectivity as a predictor of cognition in PD

A central aim of this study was to determine whether individualized MRI-based head models provide meaningful advantages over standard template models for predicting cognitive performance from EEG. The comparable predictive performance of standard template and individual head models has implications for the scalability, cost, and clinical feasibility of EEG-based biomarkers.

The strong performance of FC in the theta-, alpha-, and beta-bands aligns with prior studies showing that cognitive impairment in PD involves alterations in distributed cortico-cortical networks that manifest predominantly within these frequencies. Increases in theta power and connectivity, reductions in alpha-band synchrony, and disruptions of beta-band dynamics have been repeatedly associated with executive dysfunction, slowed processing, and global cognitive decline (Iyer et al., 2020; Gschwandtner et al., 2023; Chaturvedi et al., 2019; Sasidharan et al., 2025). Consistent with previous work linking EEG connectivity to cognitive impairment in PD (Anjum et al., 2024; Zawislak-Fornagiel et al., 2025), our findings demonstrate that these frequency-specific network features also support the prediction of global cognitive performance.

Modelling cognition as a continuous outcome allowed us to capture inter-individual variability beyond diagnostic group boundaries, which may be particularly important in PD given the gradual and heterogeneous progression of cognitive decline. In terms of magnitude, the predictive performance observed here (R² ≈ 0.07–0.17, r ≈ 0.30–0.43) falls within the range reported in broader neuroimaging-based prediction studies. Resting-state fMRI studies predicting general intelligence, for example, have reported average correlations of r ≈ 0.42 (95% CI: 0.35–0.50) (Vieira et al., 2022). In our previous work using a similar predictive framework applied to fMRI data, we observed correlations up to r ≈ 0.50 for task-based working memory and r ≈ 0.47 for resting-state fMRI (Tetereva et al., 2025). Taken together, these comparisons indicate that EEG-based connectivity measures can approach the predictive performance observed in fMRI studies, despite differences in spatial resolution and signal characteristics.

Our results show that amplitude-based FC can provide comparably robust, and in some cases numerically higher, predictive performance for cognitive outcomes in PD compared to phase-based FC measures. This result is particularly interesting given that many EEG connectivity studies (Gschwandtner et al., 2023; Chaturvedi et al., 2019; Geraedts et al., 2021; Zawislak-Fornagiel et al., 2025) tend to prioritise PLI, wPLI, or other phase-lag–based metrics. Preference for phase-based metrics reflect the common practice used to characterise inter-regional synchrony, which is not directly captured by amplitude-based measures. While phase-based measures capture fast synchronisation between neural signals, amplitude-based measures such as AEC capture slower co-fluctuations in signal power across regions that may reflect large-scale network organization and neuromodulatory influences, which are known to be affected in PD and may underpin global cognitive performance (Gratwicke et al., 2015; Slater et al., 2024). This underscores the value of including power-based (AEC) FC measures, as they provide complementary information about network dynamics relevant to cognitive function and may capture more stable large-scale connectivity patterns than phase-based metrics. This, in turn, may help explain why standard and individual head models yielded comparable predictive performance.

### Implications of similar performance between individual and standard head models

A central contribution of this study is the finding that individualized anatomical head models do not enhance prediction performance compared with standard template models. This result contrasts with simulation studies showing that template models introduce larger source localization errors (Liu et al., 2018; Vorwerk et al., 2014) and with methodological work suggesting that such errors may propagate into connectivity and ML-based analyses (Liu et al., 2023). At least for prediction of global cognitive performance using resting-state FC in PD, these theoretical benefits do not translate into measurable improvements. This highlights an important distinction between methodological accuracy (e.g., localisation error) and predictive utility.

Prior work has often focused on classification predictions and regionally constrained features, whereas the present approach uses cortex wide networks and source-space functional connectivity combined with multivariate prediction models. Several explanations may account for this. First, prediction models aggregate connectivity across the entire connectome, which may reduce sensitivity to local deviations in source geometry. In other words, although individualized head models may improve the spatial precision of dipole localization, the resulting differences may not meaningfully affect large-scale connectivity patterns once parcellated into a standardized cortical atlas comprising 360 regions. Consistent with this interpretation, feature importance analyses revealed highly similar spatial patterns between standard and individualized head models, indicating that the same networks drive predictive performance in both approaches. However, this similarity differed by the connectivity metric in that standard and individual head models showed stronger correspondence in feature-importance patterns for AEC than for dwPLI. This may suggest that AEC-based prediction is driven more strongly by within- and between-network amplitude-covariation patterns, which may be relatively robust to differences in source localisation. In contrast, dwPLI-based prediction may rely more on phase-lagged interactions distributed across multiple functional networks, and may therefore be more sensitive to localisation precision. Second, the cognitive phenotype examined here, that is, global cognitive performance aggregated across multiple cognitive domains reflects distributed brain function rather than focal neural processes. This is further supported by feature importance results showing contributions from multiple large-scale networks, including the cingulo-opercular (salience) network, the default mode network, fronto-parietal network, and somatomotor system. In particular, AEC_theta showed a clear and spatially distributed pattern of feature importance across multiple large-scale networks (Figure 3), whereas dwPLI_theta exhibited less consistent and more diffuse network-level patterns. Together, these findings support the interpretation that the predictive information was distributed across broad network-level features rather than confined to focal spatial effects. Template-induced spatial imprecision may therefore have less impact for modelling such broad phenotypes than for tasks requiring more precise localization (e.g., epilepsy source mapping). These findings have important implications for the clinical and research use of EEG-based biomarkers. Individual MRI acquisition is costly, time-consuming, and often unavailable in many clinical and even research contexts. The observation that standard head models achieve comparable performance to individualized MRI-based models suggests that EEG-based prediction pipelines can be implemented more broadly, and at lower cost, increasing the accessibility and scalability of biomarker development. This is particularly relevant for populations when routine MRI acquisition may be limited by cost, mobility constraints, or contraindications, including in community clinics and low-resource environments.

### Comparison with previous EEG–cognition prediction studies

Our findings are broadly consistent with prior work demonstrating that EEG connectivity is associated with cognitive performance in PD (Anjum et al., 2024; Betrouni et al., 2019; Chaturvedi et al., 2019; Sweeney et al., 2020; Geraedts et al., 2021; Jennings et al., 2022; Gschwandtner et al., 2023; Parajuli et al., 2023; Mostile et al., 2025; Sasidharan et al., 2025). However, most of these earlier studies used classification-based machine-learning frameworks, often reporting high accuracy values (>0.8) for distinguishing diagnostic groups such as PD-MCI and PD-N. Although such approaches are valuable for group-level discrimination, they address a different predictive task from the present study, which used regression-based prediction of continuous cognitive scores and evaluated performance using r and R². Direct comparison of performance across these approaches is therefore not straightforward. Nevertheless, both lines of evidence indicate strong associations between EEG connectivity and cognition. By modelling cognition as a continuous outcome, the present study was able to capture variability across the full spectrum of cognitive performance, including individual differences that may be obscured by categorical classification. This may be particularly relevant for clinical applications aimed at early detection or monitoring, where subtle within-subject changes may precede categorical diagnostic transitions. The present study also extends this literature by implementing source-space FC, multiple connectivity modalities, and employing a rigorous nested cross-validation framework with extensive hyperparameter tuning in a fully continuous prediction setting. Unlike the current study, previous studies that explored machine learning approaches generally relied on a limited set of algorithms (Zawislak-Fornagiel et al., 2025). In contrast, we employed multiple machine learning algorithms, enabling a more comprehensive and data-driven evaluation of predictive patterns in complex EEG data. The convergence of performance across multiple algorithms further suggests that the predictive signal is robust and not strongly dependent on a specific modelling approach.

### Limitations and future directions

The current pipeline was evaluated on eyes-closed resting-state EEG in a single sample. Nonetheless, the sample size used was moderately sized (N = 136 PD plus 51 controls) and larger than most previous studies using machine learning with EEG data in PD, which had a median size of 42 (range 33 to 118; Zawislak-Fornagiel et al., 2025). However, larger sample sizes or aggregated consortia for data harmonization may allow comparisons across sex, disease stage, and other clinical variables or post-mortem neuropathology, which may help define heterogeneity and improve prediction (Choi et al., 2024; Conti et al., 2025; Del Percio et al., 2025). In particular, modelling heterogeneity within PD, such as non-motor symptom subtypes and trajectories (Velucci et al., 2025) may represent an important next step for improving predictive accuracy. The robustness of the current findings should be evaluated in independent datasets and across different recording conditions, including eyes-open paradigms, to determine the generalizability of the predictive patterns observed here. External validation across independent cohorts will be critical to establish the clinical utility and reproducibility of these findings.

In the present study, we focused on systematically evaluating multiple individual algorithms to provide a clear and controlled comparison across modelling approaches. Future work may extend this framework by exploring ensemble or stacking machine-learning approaches, which can integrate complementary information across models and potentially improve predictive stability and performance (Tetereva et al., 2025).

Finally, interpreting the underlying neurophysiology remains an important goal. While the current analyses included initial feature importance assessments, further work is needed to systematically characterise the specific functional connections and networks driving prediction. This may provide deeper insight into the network-level alterations associated with cognitive decline in Parkinson’s disease and help guide more targeted biomarker development.

Several avenues should be pursued to build on the present findings. Restricting the delta bin to 2.5 to 4 Hz may improve convergence during model building by minimizing movement artifacts that can occur in PD samples (Caviness et al., 2007). Longitudinal work is needed to evaluate the prediction of future cognitive impairment using machine learning models of EEG activity (Geraedts et al., 2018; Choi et al., 2024). Such longitudinal prediction may be particularly valuable for identifying individuals at risk of rapid cognitive decline and informing early intervention strategies.

### Conclusions

This study demonstrates that resting-state EEG functional connectivity, particularly in the theta, alpha, and beta bands, can predict cognitive performance in Parkinson’s disease with modest but consistent accuracy. Importantly, we show that individualized MRI-based head models do not provide measurable benefits over standard templates for predicting cognition, suggesting that more accessible and computationally efficient approaches may achieve comparable performance in this context. Future work should investigate domain-specific cognitive outcomes, multimodal integration with MRI, and longitudinal applications for tracking disease subtype and the prediction of future progression. Overall, our findings support the feasibility of scalable EEG-based biomarkers of cognitive decline in PD and highlight the relevance of frequency-specific network features for prediction.

## Declarations

### Ethics approval and consent to participate

The New Zealand Parkinson’s Progression Programme (NZP3) was approved by the Northern B Health and Disability Ethics Committee of the New Zealand Ministry of Health (ethics reference: 2022 EXP 13106; approval date: 16 January 2023). The present study used data collected as part of the approved NZP3 protocol, including resting-state EEG, structural MRI, and neuropsychological assessment data. All participants provided written informed consent before participation.

### Data availability statement

The data that support the findings of this study are not publicly available due to ethical and privacy restrictions. Data may be available from the corresponding and senior authors upon reasonable request and subject to appropriate institutional and ethical approvals.

### Code availability statement

Analysis code is available at: https://github.com/HAM-lab-Otago-University/NZP3_Chch_cog_pred_STDvsIND_headmodels

### Author contributions

Alina Tetereva: Conceptualization, Methodology, Formal analysis, Investigation, Visualization, Writing – original draft, Writing – review and editing.

Grace Hall-McMaster: Data collection, Review and editing.

Nicky Slater: Data collection, Review and editing.

Abbie Harris: Data collection, Review and editing.

Reza Shoorangiz: Conceptualization, Data collection, Review and editing.

Campbell Le Heron: Review and editing.

Ross Keenan: Review and editing.

Daniel Myall: Review and editing.

Toni Pitcher: Review and editing.

Ian Kirk: Resources, Review and editing.

Wassilios G. Meissner: Review and editing.

Tim Anderson: Resources, Investigation, Review and editing.

Tracy Melzer: Resources, Investigation, Review and editing.

Narun Pat: Conceptualization, Methodology, Supervision, Writing – review and editing.

John Dalrymple-Alford: Conceptualization, Methodology, Resources, Supervision, Writing – review and editing.

## Acknowledgements

The authors thank the participants and their families for their involvement in the New Zealand Parkinson’s Progression Programme. The authors also acknowledge the research staff and clinicians involved in participant recruitment, assessment, EEG acquisition, MRI acquisition, and data management.

## Preprint statement

A previous version of this manuscript was posted as a preprint on bioRxiv: https://doi.org/10.64898/2026.05.07.723671.

## Supplementary materials

**Figure S1.**
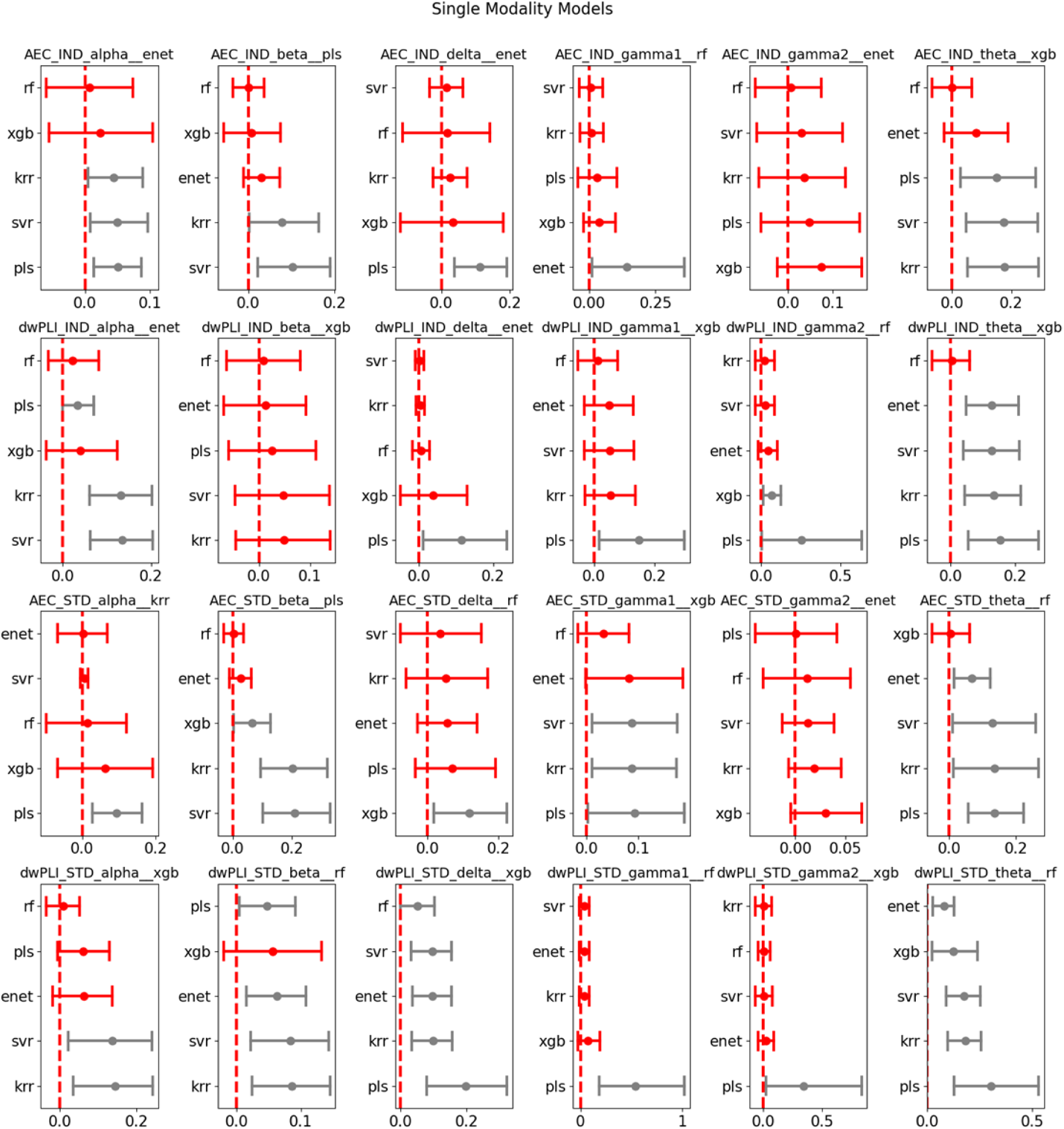
The figure shows bootstrapped difference (ΔR^2^) subplots for each modality, comparing the best-performing algorithm (i.e., the algorithm with the maximum R² value) against the performance of all other algorithms. The best-performing algorithm is indicated at the top of each subplot, while the other algorithms are listed along the y-axis. Dots represent the mean bootstrapped differences, and bars represent the 95% confidence intervals (CIs). Grey indicates a significant difference between algorithms, while red indicates no significant difference. A difference is considered significant when the 95% CI does not cross zero.

**Figure S2.**
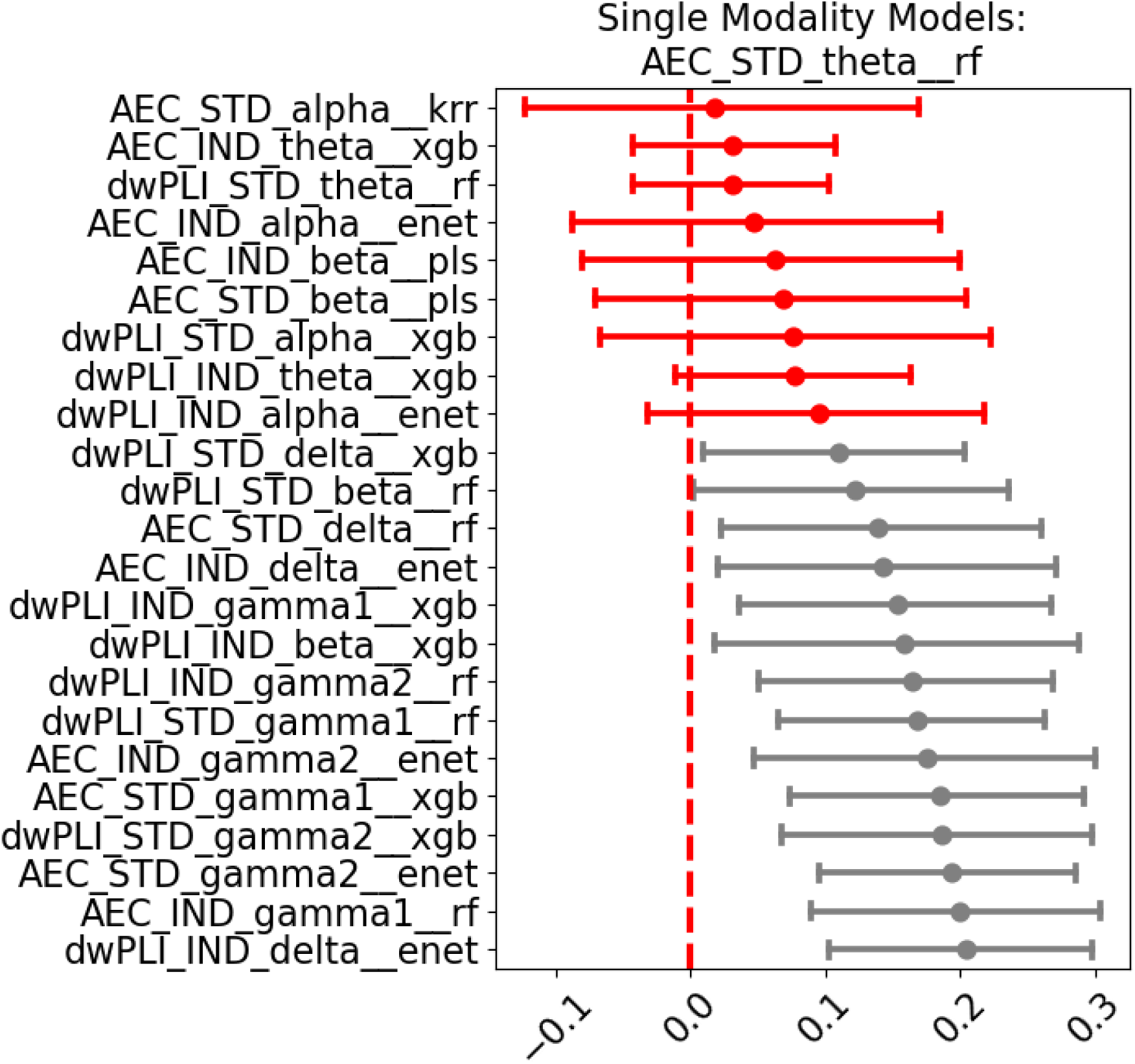
This figure shows the bootstrapped difference (ΔR^2^) comparison between the best-performing modality (using its highest-performing algorithm, indicated at the top of each subplot) and all other modalities using their respective best algorithms. The algorithms being compared are listed along the y-axis. Dots represent the mean bootstrapped differences, and bars represent the 95% confidence intervals. Grey indicates a significant difference, while red indicates no significant difference. A difference is considered significant when the 95% CI does not cross zero.

**Figure S3.**
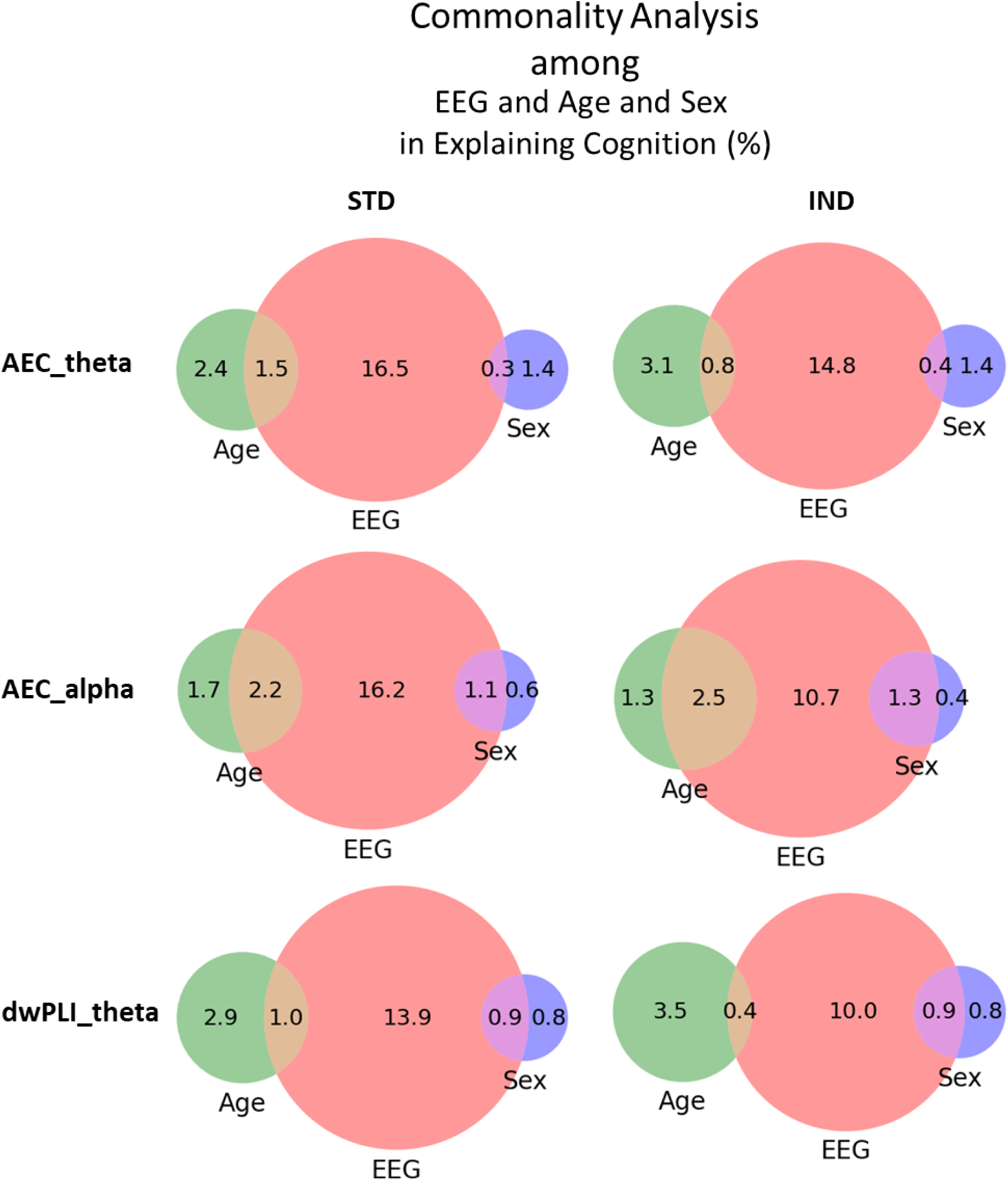
Commonality analysis comparing the variance in global cognition (global Z) explained by EEG, age, and sex in three-predictor models. Analyses are shown for the three best-performing EEG modalities: AEC theta, AEC alpha, and dwPLI theta, separately for standard (STD) and individualized (IND) head models. Commonality analysis decomposes the total variance explained by the full multiple regression model into the unique contribution of each predictor and the shared/common contribution between predictors (Nimon et al., 2008). For three regressors, this includes three unique effects, three pairwise shared effects, and one common effect shared by all three predictors. Negative common effects can occur, particularly in the presence of multicollinearity (Ray-Mukherjee et al., 2014). In the present analysis, negative common effects were observed for the shared Age–Sex component and the shared EEG–Age–Sex component. These negative values were treated as zero (Frederick, 1999), and the remaining effects were scaled proportionally to the total variance explained by the full regression model.

## References

Aarsland, D., Batzu, L., Halliday, G. M., et al. (2021). Parkinson disease–associated cognitive impairment. Nature Reviews Disease Primers, 7(1), 1–21.

Abreu, R., Simões, M., & Castelo-Branco, M. (2020). Pushing the limits of EEG: Estimation of large-scale functional brain networks and their dynamics validated by simultaneous fMRI. Frontiers in Neuroscience. 10.3389/fnins.2020.00423

Almgren, H., Camacho, M., Hanganu, A., Kibreab, M., Camicioli, R., Ismail, Z., et al. (2023). Machine learning-based prediction of longitudinal cognitive decline in early Parkinson’s disease using multimodal features. Scientific Reports, 13(1), 13193.

Anjum, M. F., Espinoza, A. I., Cole, R. C., Singh, A., May, P., Uc, E. Y., et al. (2024). Resting-state EEG measures cognitive impairment in Parkinson’s disease. npj Parkinson’s Disease, 10, Article 6. 10.1038/s41531-023-00598-5

Appelhoff, S., Sanderson, M., Brooks, T. L., Vliet, M., Quentin, R., Holdgraf, C., et al. (2019). MNE-BIDS: Organizing electrophysiological data into the BIDS format and facilitating their analysis. Journal of Open Source Software, 4(44), 1896. 10.21105/joss.01896

Babiloni, C., Arakaki, X., Azami, H., Bennys, K., Blinowska, K., Bonanni, L., et al. (2021). Measures of resting state EEG rhythms for clinical trials in Alzheimer’s disease: Recommendations of an expert panel. Alzheimer’s & Dementia, 17(9), 1528–1553.

Babiloni, C., Del Percio, C., Lizio, R., Noce, G., Lopez, S., Soricelli, A., et al. (2018a). Abnormalities of resting-state functional cortical connectivity in patients with dementia due to Alzheimer’s and Lewy body diseases: An EEG study. Neurobiology of Aging, 65, 18–40.

Babiloni, C., Del Percio, C., Lizio, R., Noce, G., Lopez, S., Soricelli, A., et al. (2018b). Functional cortical source connectivity of resting state electroencephalographic alpha rhythms shows similar abnormalities in patients with mild cognitive impairment due to Alzheimer’s and Parkinson’s diseases. Clinical Neurophysiology, 129(4), 766–782.

Bénar, C. G., & Gotman, J. (2002). Modeling of post-surgical brain and skull defects in the EEG inverse problem with the boundary element method. Clinical Neurophysiology, 113(1), 48–56. 10.1016/S1388-2457(01)00714-3

Betrouni, N., Delval, A., Chaton, L., et al. (2019). Electroencephalography-based machine learning for cognitive profiling in Parkinson’s disease: Preliminary results. Movement Disorders, 34(2), 210–217.

Bigdely-Shamlo, N., Mullen, T., Kothe, C., Su, K.-M., & Robbins, K. A. (2015). The PREP pipeline: Standardized preprocessing for large-scale EEG analysis. Frontiers in Neuroinformatics, 9, 16. 10.3389/fninf.2015.00016

Breiman, L. (2001). Random forests. Machine Learning, 45(1), 5–32. 10.1023/A:1010933404324

Brydges, C. R. (2019). Effect size guidelines, sample size calculations, and statistical power in gerontology. Innovation in Aging, 3(4), 1–8.

Burchill, E., Watson, C. J., Fanshawe, J. B., Badenoch, J. B., Rengasamy, E., Ghanem, D. A., et al. (2024). The impact of psychiatric comorbidity on Parkinson’s disease outcomes: A systematic review and meta-analysis. The Lancet Regional Health – Europe, 39.

Caviness, J. N., Hentz, J. G., Evidente, V. G., Driver-Dunckley, E., Samanta, J., Mahant, P., et al. (2007). Both early and late cognitive dysfunction affects the electroencephalogram in Parkinson’s disease. Parkinsonism & Related Disorders, 13(6), 348–354.

Chaturvedi, M., Bogaarts, J. G., Kozak, V. V., et al. (2019). Phase lag index and spectral power as QEEG features for identification of patients with mild cognitive impairment in Parkinson’s disease. Clinical Neurophysiology, 130(10), 1937–1944.

Chen, T., & Guestrin, C. (2016). XGBoost: A scalable tree boosting system. In Proceedings of the 22nd ACM SIGKDD International Conference on Knowledge Discovery and Data Mining (pp. 785–794). 10.1145/2939672.2939785

Choi, A., Zhang, N., Adler, C. H., Beach, T. G., Shill, H. A., Driver-Dunckley, E., et al. (2024). Resting-state EEG predicts cognitive decline in a neuropathologically diagnosed longitudinal community autopsied cohort. Parkinsonism & Related Disorders, 128, 107120.

Conti, M., Bovenzi, R., Pierantozzi, M., Simonetta, C., Ferrari, V., Bissacco, J., et al. (2025). Sex hormones shape EEG-based functional connectivity in early-stage Parkinson’s disease patients. NeuroImage: Clinical, 45, 103721.

de Cheveigné, A., & Simon, J. Z. (2008). Denoising based on time-shift PCA. Journal of Neuroscience Methods, 165(2), 297–305.

Del Percio, C., Lizio, R., Lopez, S., Noce, G., Jakhar, D., Carpi, M., et al. (2025). Resting-state electroencephalographic rhythms depend on sex in patients with dementia due to Parkinson’s and Lewy body diseases. Neurobiology of Disease, 206, 106807.

Dierks, T., Jelic, V., Pascual-Marqui, R., Wahlund, L., Julin, P., Linden, D., et al. (2000). Spatial pattern of cerebral glucose metabolism correlates with localization of intracerebral EEG generators in Alzheimer’s disease. Clinical Neurophysiology, 111(10), 1817–1824.

Droby, A., Nosatzki, S., Edry, Y., et al. (2022). The interplay between structural and functional connectivity in early-stage Parkinson’s disease. Journal of the Neurological Sciences, 442, 120452.

Drucker, H., Burges, C. J. C., Kaufman, L., Smola, A., & Vapnik, V. (1996). Support vector regression machines. In Advances in Neural Information Processing Systems.

Efron, B., & Tibshirani, R. J. (1994). An introduction to the bootstrap. Chapman & Hall/CRC.

Emre, M., Aarsland, D., Brown, R., Burn, D. J., Duyckaerts, C., Mizuno, Y.,… & Dubois, B. (2007). Clinical diagnostic criteria for dementia associated with Parkinson’s disease. Movement disorders: official journal of the Movement Disorder Society, 22(12), 1689–1707.

Fischl, B. (2012). FreeSurfer. NeuroImage, 62(2), 774–781.

Fischl, B., Sereno, M. I., Tootell, R. B. H., & Dale, A. M. (1999). High-resolution intersubject averaging and a coordinate system for the cortical surface. Human Brain Mapping, 8(4), 272–284.

Fu, Y., & Halliday, G. M. (2025). Dementia with Lewy bodies and Parkinson disease dementia—the same or different and is it important? Nature Reviews Neurology.

Fuchs, M., Wagner, M., & Kastner, J. (2001). Boundary element method volume conductor models for EEG source reconstruction. Clinical Neurophysiology, 112(8), 1400–1407.

Funder, D. C., & Ozer, D. J. (2019). Evaluating effect size in psychological research: Sense and nonsense. Advances in methods and practices in psychological science, 2(2), 156–168.

Gallagher, J., Gochanour, C., Caspell-Garcia, C., Dobkin, R. D., Aarsland, D., Alcalay, R. N., et al. (2024). Long-term dementia risk in Parkinson disease. Neurology, 103(5), e209699.

Geraedts, V. J., Boon, L. I., Marinus, J., Gouw, A. A., van Hilten, J. J., Stam, C. J., et al. (2018). Clinical correlates of quantitative EEG in Parkinson disease: A systematic review. Neurology, 91(19), 871–883.

Geraedts, V. J., Koch, M., Contarino, M. F., et al. (2021). Machine learning for automated EEG-based biomarkers of cognitive impairment during deep brain stimulation screening in patients with Parkinson’s disease. Clinical Neurophysiology, 132(5), 1041–1048.

Gignac, G. E., & Szodorai, E. T. (2016). Effect size guidelines for individual differences researchers. Personality and individual differences, 102, 74–78.

Glasser, M. F., Coalson, T. S., Robinson, E. C., Hacker, C. D., Harwell, J., Yacoub, E., et al. (2016). A multi-modal parcellation of human cerebral cortex. Nature, 536(7615), 171–178.

Gratwicke, J., Jahanshahi, M., & Foltynie, T. (2015). Parkinson’s disease dementia: a neural networks perspective. Brain, 138(6), 1454–1476.

Gschwandtner, U., Bogaarts, G., Roth, V., & Fuhr, P. (2023). Prediction of cognitive decline in Parkinson’s disease patients with EEG connectivity characterized by time-between-phase-crossing. Scientific Reports, 13(1), 5093.

Guo, X., et al. (2025). Research progress of multimodal biomarkers in the early diagnosis of mild cognitive impairment in Parkinson’s disease. Frontiers in Neurology, 16, 1652378.

Hallez, H., Vanrumste, B., Van Hese, P., Delputte, S., & Lemahieu, I. (2008). Dipole estimation errors due to anisotropic conductivity modelling. Physics in Medicine and Biology, 53(7), 1877–1894.

Haufe, S., Meinecke, F., Görgen, K., Dähne, S., Haynes, J.-D., Blankertz, B., & Bießmann, F. (2014). On the interpretation of weight vectors of linear models in multivariate neuroimaging. NeuroImage, 87, 96–110.

Horne, K. L., MacAskill, M. R., Myall, D. J., Livingston, L., Grenfell, S., Pascoe, M. J., et al. (2021). Neuropsychiatric symptoms and dementia in Parkinson’s disease. Movement Disorders Clinical Practice, 8(3), 390–399.

Iyer, K. K., Au, T. R., Angwin, A. J., Copland, D. A., & Dissanayaka, N. N. W. (2020). Theta and gamma connectivity is linked with affective and cognitive symptoms in Parkinson’s disease. Journal of Affective Disorders, 277, 875–884.

Jennings, J. L., Peraza, L. R., Baker, M., et al. (2022). EEG for assisting dementia diagnosis. Alzheimer’s Research & Therapy, 14(1), 109.

Ji, J. L., Spronk, M., Kulkarni, K., Repovš, G., Anticevic, A., & Cole, M. W. (2019). Mapping the human brain’s cortical-subcortical functional network organization. NeuroImage, 185, 35–57.

Lawson, R. A., Yarnall, A. J., Duncan, G. W., et al. (2016). Cognitive decline and quality of life in Parkinson’s disease. Parkinsonism & Related Disorders, 27, 47–53.

Lawson, R. A., Yarnall, A. J., Johnston, F., et al. (2017). Cognitive impairment in Parkinson’s disease: Impact on quality of life of carers. International Journal of Geriatric Psychiatry, 32(12), 1362–1370.

Lee, D. G., & Lee, S. B. (2025). EEG feature extraction for neurodegenerative disease diagnosis. Digital Health Research, 3(3).

Litvan, I., Goldman, J. G., Tröster, A. I., et al. (2012). Diagnostic criteria for mild cognitive impairment in Parkinson’s disease. Movement Disorders, 27(3), 349–356.

Liu, C., Downey, R. J., Mu, Y., Richer, N., Hwang, J., Shah, V. A., et al. (2023). Comparison of EEG source localization using simplified and anatomically accurate head models. IEEE Transactions on Neural Systems and Rehabilitation Engineering, 31, 2591–2602.

Liu, Q., Ganzetti, M., Wenderoth, N., & Mantini, D. (2018). Detecting large-scale brain networks using EEG. Frontiers in Neuroinformatics, 12, Article 4.

MacAskill, M. R., Pitcher, T. L., Melzer, T. R., Myall, D. J., Horne, K. L., Shoorangiz, R., et al. (2022). The New Zealand Parkinson’s progression programme. Journal of the Royal Society of New Zealand, 53(4), 466–488.

Mahajan, A., Jaramillo-Jimenez, A., D’Anselmo, A., Prete, G., Bristot, L., Varanese, S., et al. (2025). Quantitative EEG in dementia with Lewy bodies. Clinical Neurophysiology Practice, 10, 222–233.

Michel, C. M., Brunet, D., Murray, M. M., & Koenig, T. (2019). EEG source imaging: A practical review. Frontiers in Neurology, 10, 325.

MNE-Connectivity Developers. (2024). mne-connectivity (Version 0.7.0).

Mostile, G., Quattropani, S., Contrafatto, F., et al. (2025). Machine learning for cognitive profiles in Parkinson’s disease. Computational and Structural Biotechnology Journal.

Nimon, K., Lewis, M., Kane, R., & Haynes, R. M. (2017). Erratum to: An R package to compute commonality coefficients in the multiple regression case: An introduction to the package and a practical example. Behavior Research Methods, 49, 2275.

Parajuli, M., Amara, A. W., & Shaban, M. (2023). Screening of mild cognitive impairment using deep learning. In IEEE EMBS Neural Engineering Conference.

Pedregosa, F., Varoquaux, G., Gramfort, A., et al. (2011). Scikit-learn: Machine learning in Python. Journal of Machine Learning Research, 12, 2825–2830.

Pion-Tonachini, L., Kreutz-Delgado, K., & Makeig, S. (2019). ICLabel: EEG independent component classifier. NeuroImage, 198, 181–197.

Ramon, C., Schimpf, P. H., & Haueisen, J. (2006). Influence of head models on EEG source localization. Biomedical Engineering Online, 5, Article 10.

Sasidharan, D., Sowmya, V., & Gopalakrishnan, E. A. (2025). EEG complexity analysis for Parkinson’s disease classification. Physical and Engineering Sciences in Medicine.

Saunders, C., Gammerman, A., & Vovk, V. (1998). Ridge regression learning algorithm. In International Conference on Machine Learning.

Schapira, A. H. V., Chaudhuri, K. R., & Jenner, P. (2017). Non-motor features of Parkinson disease. Nature Reviews Neuroscience, 18(7), 435–450.

Slater, N. M., Melzer, T. R., Myall, D. J., Anderson, T. J., & Dalrymple-Alford, J. C. (2024). Cholinergic basal forebrain integrity and cognition in Parkinson’s disease: a reappraisal of magnetic resonance imaging evidence. Movement disorders, 39(12), 2155–2172.

Spoa, M., Monti, S., Bjekić, J., Guerra, A., Fiorenzato, E., Cauzzo, S., et al. (2025). Resting-state EEG changes in Parkinson’s disease. Journal of Neural Transmission.

Svenningsson, P., Westman, E., Ballard, C., & Aarsland, D. (2012). Cognitive impairment in Parkinson’s disease. The Lancet Neurology, 11(8), 697–707.

Sweeney, A., Devereux, B., Ong, C., et al. (2020). EEG and machine learning in Parkinson’s disease. Alzheimer’s & Dementia, 16, e040432.

Tetereva, A., Knodt, A. R., Melzer, T. R., van der Vliet, W., Gibson, B., Hariri, A. R., et al. (2022). Capturing brain–cognition relationships. NeuroImage, 265, 119737.

Tetereva, A., Knodt, A. R., Melzer, T. R., van der Vliet, W., Gibson, B., Hariri, A. R., et al. (2025). Improving predictability of brain–cognition associations. PNAS Nexus, 4(6), pgaf175.

Vanrumste, B., Van Hoey, G., Van de Walle, R., D’Havé, M., Lemahieu, I., & Boon, P. (2000). Dipole location errors in EEG source analysis. Medical & Biological Engineering & Computing, 38(5), 528–534.

Velucci, V., Iliceto, G., Vitucci, B., Idrissi, S., Milella, G., Mascia, M. M.,… & Parkinson’s Progression Markers Initiative. (2025). Non-motor symptom subtypes in early Parkinson’s disease. Parkinsonism & Related Disorders, 107982.

Vieira, B. H., Pamplona, G. S. P., Fachinello, K., Silva, A. K., Foss, M. P., & Salmon, C. E. G. (2022). On the prediction of human intelligence from neuroimaging: A systematic review of methods and reporting. Intelligence, 93, 101654. DOI: 10.1016/j.intell.2022.101654

Vorwerk, J., Cho, J.-H., Rampp, S., Hamer, H., Knösche, T. R., & Wolters, C. H. (2014). Head volume conductor modeling in EEG and MEG. NeuroImage, 100, 590–607.

Weintraub, D., Aarsland, D., Biundo, R., Dobkin, R., Goldman, J., & Lewis, S. (2022). Management of psychiatric and cognitive complications in Parkinson’s disease. BMJ, 379.

Wold, S., Sjöström, M., & Eriksson, L. (2001). PLS regression. Chemometrics and Intelligent Laboratory Systems, 58(2), 109–130.

Wood, K. L., Myall, D. J., Livingston, L., Melzer, T. R., Pitcher, T. L., MacAskill, M. R., et al. (2016). PD-MCI criteria and dementia risk. npj Parkinson’s Disease, 2, 1–8.

Zawiślak-Fornagiel, K., Gorzkowska, A., Galus, W., Węgrzynek-Gallina, J., Ledwoń, D., & Mitas, A. W. (2025). Quantitative EEG and machine learning in PD. Clinical Neurophysiology.

Zou, H., & Hastie, T. (2005). Regularization and variable selection via elastic net. Journal of the Royal Statistical Society: Series B, 67(2), 301–320.

## References

Frederick, B. N. (1999). Partitioning variance in the multivariate case: A step-by-step guide to canonical commonality analysis. Advances in Social Science Methodology, 5, 305–318.

Ray-Mukherjee, J., Nimon, K., Mukherjee, S., Morris, D. W., Slotow, R., & Hamer, M. (2014). Using commonality analysis in multiple regressions: A tool to decompose regression effects in the face of multicollinearity. Methods in Ecology and Evolution, 5, 320–328.

